# Adolescent Stress Increases Adult Ethanol Self-administration and Alters Ventral Tegmental Area GABA Signaling

**DOI:** 10.1101/715607

**Authors:** David A Connor, Ruthie E Wittenberg, Jillian Drogin, Allison Mak, John A Dani

## Abstract

Alcohol use disorders (AUDs) continue to be a significant public health problem. Early life stress and adversity have long-lasting effects on a wide range of behaviors, including responses to drugs of abuse. Epidemiological evidence indicates that exposure to early life stress contributes to alcohol use disorders and, while it is known that stress and alcohol both act on overlapping mesolimbic circuitry, the cellular mechanisms underlying the relationship between stress and alcohol intake are not well understood. Previous work has demonstrated that acute stress increases ethanol intake mediated by changes in GABA signaling within the ventral tegmental area (VTA). Here we investigated if adolescent stress exposure might elicit long-term, persistent increases in ethanol self-administration associated with altered VTA GABA signaling. To this end, we exposed adolescent postnatal day (PND) 28 male rats to 14 days of chronic variable stress (CVS) and then examined operant ethanol self-administration begun at least 30 days later. We found that adolescent stress exposure resulted in significantly increased ethanol self-administration in adulthood. In contrast, adult (PND 82) male rats exposed to the same CVS protocol did not display increased ethanol self-administration that was begun 30 days later. Furthermore, we found that adolescent stress exposure resulted in enhancement of ethanol-induced GABA signaling onto VTA dopamine neurons and impairments in VTA GABA chloride homeostasis. The results indicate that adolescence is a period vulnerable to stress, which produces long-term changes in VTA GABA signaling associated with increased ethanol self-administration behavior.

## Introduction

Alcohol use disorders (AUDs) are a significant public health problem, with an estimated 88,000 alcohol-related deaths annually in the United States (Stahre, Roeber, Kanny, Brewer, & Zhang, 2014). Stress is an important risk factor for the development of substance use disorders (Uhart & Wand, 2009). In humans, early life trauma and stress predict earlier onset of drinking and increased alcohol intake, thereby, facilitating development of AUDs (Dawson & Archer, 1993; Enoch, 2011; Kaufman et al., 2007). Preclinical rodent models demonstrate that acute stress during adulthood increases ethanol seeking and self-administration (Le et al., 1998; Ostroumov et al., 2016). Additionally, early life stress has been shown to increase later ethanol intake (Butler, Karkhanis, Jones, & Weiner, 2016; Lopez, Doremus-Fitzwater, & Becker, 2011).

Adolescence represents a time of critical neurodevelopment, including maturation of the neural circuitry and neurotransmitter systems (Spear, 2000; Steinberg, 2005). As a result, such developing neural systems of adolescent individuals are vulnerable to long-lasting consequences of perturbations, such as stress. Herein, we employ chronic variable stress (CVS), a non-habituating chronic stress paradigm (Herman, 2013), to examine the long-term effects of adolescent stress on adult changes in VTA neurophysiology and the associated ethanol self-administration. We selected chronic variable and unpredictable stress exposure because it models chronic stress in rats, where it has been shown to modulate the response to drugs of abuse, including alcohol (Lopez et al., 2011). We sought to examine long-term stress-induced changes in behavior and neurophysiology isolated from the proximal effects of stress, such as immediate fear and behavioral freezing. Therefore, CVS allowed for stress exposure only during adolescence (PND 28-42), after which the stressed animals were treated similar to controls in a stress-free home environment. Also, because social isolation is a known stressor (Gentsch, Lichtsteiner, & Feer, 1981; Hatch et al., 1965), we sought to further reduce the influence of proximal stress during ethanol intake measurements by using a timed operant paradigm. In contrast to free choice ethanol intake assays, which depend on single housing, operant self-administration allowed for group housing throughout the study.

The effects of stress on alcohol intake are mediated, in part, by alterations of reward circuitry (Ostroumov & Dani, 2018a; Spanagel, Noori, & Heilig, 2014). In particular, inhibitory GABA signaling within the VTA has been shown to be an important modulator of ethanol self-administration (Doyon et al., 2013; Ostroumov & Dani, 2018a; Stobbs et al., 2004). Previous work from our lab demonstrated that acute stress in adult rats enhances ethanol-induced GABA release onto DA neurons due to adaptations in local VTA GABA neurons (Ostroumov et al., 2016). Normal functioning of GABA_A_ receptor inhibitory signaling is dependent on the extrusion of intracellular Cl^−^ (Hewitt, Wamsteeker, Kurz, & Bains, 2009). Importantly, acute adult stress was shown to decrease Cl^−^ extrusion capacity of VTA GABA neurons causing hyper-excitability of local GABAergic neurons, which increased ethanol induced inhibition of VTA DA neurons (Ostroumov et al., 2016). While these stress-induced changes have been shown to increase alcohol intake in a model of adult acute stress, it has yet to be investigated in a translational model of early-life adolescent stress. Furthermore, we aimed to determine whether chronic stress applied during adolescence produced long-lasting effects that would impact adult alcohol self-administration, as suggested by adolescent vulnerability in humans (Enoch, 2011; Pilowsky, Keyes, & Hasin, 2009).

We observed that when rats were exposed to stress during adolescence, but not during adulthood, they showed a long-term increase in ethanol self-administration. Due to the prolonged behavioral impact of stress exposure during adolescence, we hypothesized that altered VTA GABA transmission in response to ethanol would be observed later in adulthood as a consequence of adolescent stress. In a separate cohort, we investigated if adolescent stress resulted in long-term changes in GABA_A_ receptor inputs to VTA DA neurons. We found that adolescent stress did increase ethanol-induced inhibition of VTA DA neurons. Also, we examined if adolescent stress disrupted Cl^−^ homeostasis in VTA GABA neurons, and observed diminished Cl^−^ transport function in adulthood.

## Methods

### Subjects

Male Long-Evans rats (Harlan-Envigo) were received with dam at PND 20 for adolescent stress exposure or PND 75 for adult stress exposure. All rats were allowed to acclimate to the vivarium for 7 days. Adolescent rats were received in groups of 4-6, weaned at PND 21-25, and split into 2 age-matched littermate groups for stress and control groups. Similarly, adult rats were shipped with littermates and divided into equal sized age-matched stress and control groups. To minimize social stress, rats were housed by experimental group (stress or control) continuously throughout the entirety of the study. All rats were maintained in a quiet, temperature-and humidity-controlled satellite facility under a 12-hour light/dark cycle. Rats had food and water available ad libitum in their home cage, as well as similar enrichment (translucent tunnel). All procedures were carried out in compliance with guidelines specified by the Institutional Animal Care and Use Committee at University of Pennsylvania.

### Operant ethanol self-administration

Standard two-lever operant chambers with motorized spout (Med Associates Inc., St. Albans, VT, USA), were used for the self-administration experiments. All rats were handled daily (10 min) at least 5 days prior to the onset of behavioral procedures. Self-administration procedures were based on prior work from our lab (Doyon et al., 2013; Ostroumov et al., 2016; Thomas et al., 2018). Briefly, behavioral shaping occurred in 60 minute daily sessions during which both levers were active and rats had access to saccharin solution (0.1%, w/v) on a fixed ratio 1 (FR1) schedule. After initial shaping and magazine training, for all subsequent sessions, one lever functioned as the active lever and completion of the FR1 schedule elicited a cue light and access to the sipper for 10 seconds. Lever presses on the inactive lever had no function, but were recorded. Baseline saccharin intake was monitored and was measured over multiple days until responding was stable (< 25% variation) for at least 2 consecutive days. Rats were excluded if their saccharin intake stabilized above 15 ml or below 5 ml. When saccharin responding stabilized, ethanol self-administration commenced. For acquisition of ethanol self-administration, ethanol was introduced into the saccharin drinking solution in the following way: 2% ethanol on day 1 and 4% ethanol for 7 days. After acquisition, saccharin was faded out of the solution and alcohol was increased to 10% over 8 days. Finally, we measured self-administration of unsweetened 10% alcohol for 10 days.

### Chronic Variable Stress

Chronic variable stress was administered to subjects over 14 days (Table 1): adolescent, PND 28-42; adult, PND 82-96. Control rats were housed identically to stressed rats and were exposed to comparable daily handling/weighing, but with no stress exposure. Stress treated rats were exposed to 4 unique 1 hour day-time stressors, 1 or 2 per day during the light period, from the following list: restraint stress,1 hour of immobilization in a clear cylindrical Broome-style restrainer; rocker motion, individually placed in empty cage with no bedding atop a VariMix rocker set to 30 RPM; elevated platform, individually placed on a 28 cm circular platform 1 m above the floor; predator odor, Tink’s Red Fox-P is sprayed onto gauze and placed in a tube with small air holes and rats are individually exposed to the odor in an empty cage with no bedding. These daily stressors were applied such that each stressor was administered 4 times over the course of 14 days non-consecutively. Additionally, twice throughout the 14 days, rats were exposed to overnight wet bedding. For overnight wet bedding, approximately 750 ml of tap water is applied to home cage bedding and enrichment tube was removed overnight.

**Table 1.**
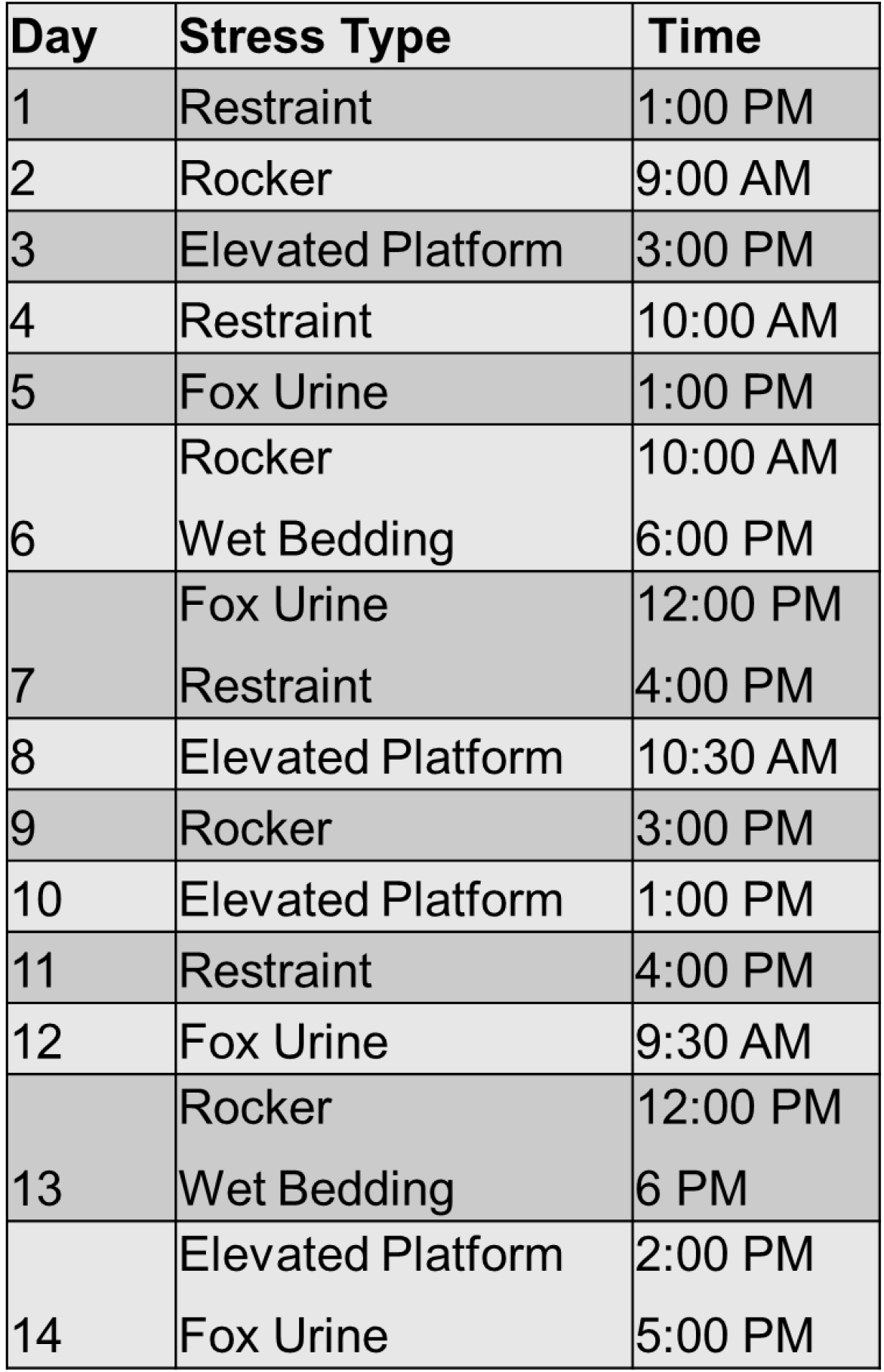
Schedule of chronic variable stress procedure over 14 days.

| **Day** | **Stress Type** | **Time** |
| --- | --- | --- |
| 1 | Restraint | 1:00 PM |
| 2 | Rocker | 9:00 AM |
| 3 | Elevated Platform | 3:00 PM |
| 4 | Restraint | 10:00 AM |
| 5 | Fox Urine | 1:00 PM |
| 6 | Rocker | 10:00 AM |
|  | Wet Bedding | 6:00 PM |
| 7 | Fox Urine | 12:00 PM |
|  | Restraint | 4:00 PM |
| 8 | Elevated Platform | 10:30 AM |
| 9 | Rocker | 3:00 PM |
| 10 | Elevated Platform | 1:00 PM |
| 11 | Restraint | 4:00 PM |
| 12 | Fox Urine | 9:30 AM |
| 13 | Rocker | 12:00 PM |
|  | Wet Bedding | 6 PM |
| 14 | Elevated Platform | 2:00 PM |
|  | Fox Urine | 5:00 PM |

### Ex Vivo Electrophysiology

Horizontal slices (230 μm) containing the VTA were cut (Leica Microsystems) from adult Long-Evans rats in ice-cold, oxygenated (95% O2, 5% CO2), high-sucrose artificial cerebrospinal fluid (ACSF) (in millimolar): 205.0 sucrose, 2.5 KCl, 21.4 NaHCO3, 1.2 NaH2PO4, 0.5 CaCl2, 7.5 MgCl2, and 11.1 dextrose. After cutting, slices were immediately transferred to normal ACSF buffer, which contained (in millimolar): 120.0 NaCl, 3.3 KCl, 25.0 NaHCO3, 1.2 NaH2PO4, 2.0 CaCl2, 1.0 MgCl2, 10.0 dextrose, and 20.0 sucrose. The slices were constantly oxygenated (95% O2, 5%CO2) and maintained at 32°C in ACSF for 40 min, then at room temperature for at least one hour. For electrophysiological recordings, slices were transferred to a holding chamber and perfused with normal ACSF at a constant rate of 2–3 mL/min at 32°C. Patch electrodes were made of thin-walled borosilicate glass with an inner diameter of 1.12 mm and an outer diameter of 1.5 mm (World Precision Instruments [WPI]) and had resistances of 1.0–2.0 MΩ when filled with the internal solution.

Spontaneous inhibitory postsynaptic currents (sIPSCs) in VTA dopamine neurons were recorded in voltage-clamp mode at −60 mV in whole-cell configuration. The internal solution used for this experiment contained (in millimolar): 135.0 KCl, 12.0 NaCl, 2.0 Mg-ATP, 0.5 EGTA, 10.0 HEPES, and 0.3 Tris-GTP (pH 7.2–7.3). During the experiment, synaptic GABA_A_ receptor currents were isolated pharmacologically with AMPA and NMDA-type glutamate receptor antagonists: 6,7 dinitroquinoxaline-2,3-dione (DNQX, 20 μM; Sigma Aldrich) and DL-2-amino-5-phosphonopentanoic acid (AP5, 50 μM; Tocris Bioscience). Data files were recorded in two-minute increments in order to establish the baseline recording before bath application of ethanol. Once a stable baseline was achieved, ethanol was washed onto the slice and recorded in two-minute increments. The baseline file immediately preceding ethanol application was then analyzed and compared to the stable ethanol file. Both the amplitude and frequency of events were measured

To measure activity-dependent depression of evoked IPSCs, whole-cell recordings were performed in VTA GABA neurons during repetitive stimulation (Hewitt et al., 2009). The internal solution contained (in milllimolar): 123.0 K+-gluconate, 8.0 NaCl, 2.0 Mg-ATP, 0.2 EGTA, 10.0 HEPES, and 0.3 Tris-GTP (pH 7.2–7.3). Throughout the experiment, synaptic GABAA receptor inputs were isolated with DNQX, AP5, and CGP55845 (1 μM, Sigma Aldrich). The liquid junction potential between the bath and the pipette solutions was corrected during recordings.

### Lateral VTA Neuron Identification

VTA dopamine neurons in the lateral VTA were identified by several factors previously used by our lab and others (Ostroumov et al., 2016), including their relatively large somata size (>20 μm), low firing frequency (<5 Hz), and the presence of a large h current (I_h_). GABA neurons were identified in the lateral VTA by their small somata size (<20 μm), high firing rate (>7 Hz), and lack of I_h_.

### Data analysis

For adolescent and adult CVS self-administration studies, data were analyzed using mixed-design ANOVA (Session Day x Stress Condition) and independent samples *t*-tests (SPSS 16.0). An independent samples *t*-test was used to assess differences between the mean sIPSC frequencies and amplitudes. For analysis of repetitive synaptic stimulation, an exponential function was fit to the group, binned datasets and compared with an extra sum of squares F-test. These statistical analyses were performed in Origin. Significance for all experiments was determined by *p* < 0.05.

## Results

### Adolescent stress increases adult ethanol self-administration

To test if adolescent stress results in increased ethanol self-administration later in adulthood, adolescent rats were exposed to 14 days of CVS starting on PND 28 (Fig. 1A). Thirty days after stress cessation, rats were handled, trained to self-administer saccharin, and then self-administer ethanol at greater than PND 72. Once rats were trained to reliably self-administer saccharin, and saccharin responding was stable, ethanol was introduced into the saccharin sweetened solution (2%-4%). Saccharin intake across the 2 days immediately prior to introduction of ethanol did not differ between adolescent control (Mean = 9.7 ml, SD = 2.5 ml, n = 12) and adolescent stress groups (Mean = 11.4 ml, SD = 2.3 ml, n = 8), *p* > 0.05.

**Figure 1.**
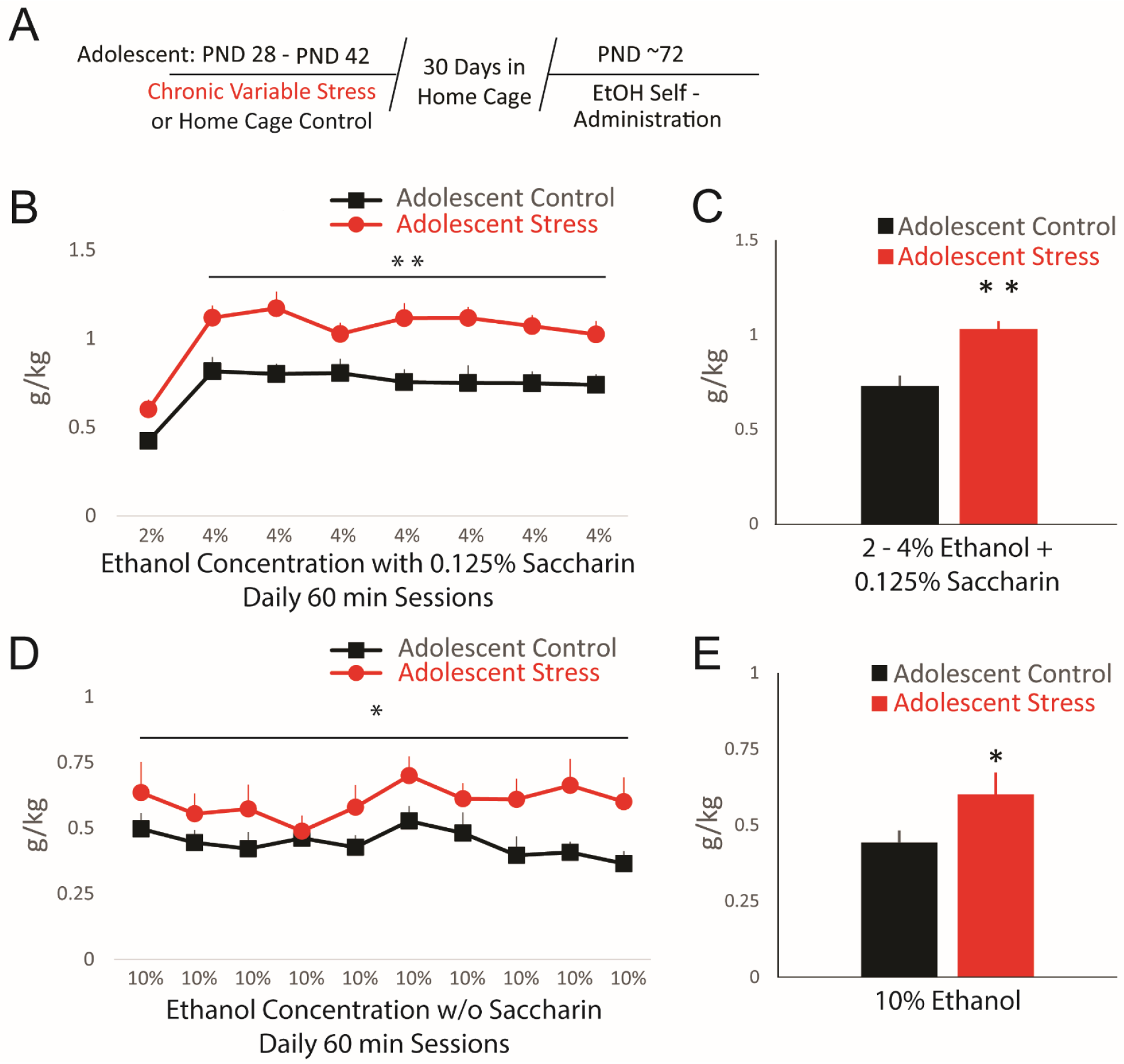
(A) Adolescent rats were exposed to chronic variable stress for 14 days, PND 28 - 42. Ethanol self-administration was assessed at least 30 days after cessation of stress. (B) Mean daily ethanol intake during initial self-administration sessions. Adolescent stress-exposed rats (red) showed greater daily intake of sweetened (0.1% saccharin) 4% ethanol intake compared to home cage controls (black). \*\**p* < 0.01, significant between subjects main effect, mixed-design ANOVA, n = 8, 11 rats per group. (C) Average ethanol self-administration of 2-4% ethanol with saccharin 0.1%. Adolescent stress exposed rats showed greater ethanol intake compared to controls. \*\**p* < 0.01, independent samples t-test, n = 8, 11 rats per group. (D) Mean daily ethanol intake during 10 daily self-administration sessions. Adolescent stress-treated rats showed greater daily intake of unsweetened 10% ethanol compared to home cage controls. \**p* < 0.05, significant between subjects main effect, mixed-design ANOVA, n = 8, 11 rats per group. (E) Average ethanol self-administration of unsweetened 10% ethanol over 10 sessions. Adolescent stress exposed rats showed greater ethanol intake compared to controls. \**p* < 0.05, independent samples t-test, n = 8, 11 rats per group.

After saccharin self-administration was stable, ethanol was introduced into the saccharin solution. Chronic variable stress during adolescence resulted in increased self-administration of sweetened ethanol solution (4.0% EtOH + 0.1% Sac) during 7 days, F(1, 17) = 19.06, *p* < 0.01; n = 8 – 11 (Fig.1B) in adulthood. Mean ethanol dose across 8 days of sweetened 2-4% ethanol was 0.73 g/kg (SD = 0.17 g/kg) for control rats and 1.03 g/kg (SD = 0.11 g/kg) for adolescent stress rats, t(17) = 4.33, *p* < 0.01 (Fig.1C). After fading saccharin out of the drinking solution and increasing alcohol to 10%, adolescent stress exposed rats showed increased unsweetened ethanol self-administration across 10 days, F(1,17) = 4.87, *p* < 0.05; n = 8 – 11 (Fig. 1D). Mean self-administered dose for adolescent stress exposed rats over 10 days of 10% ethanol was 0.6 g/kg, (SD = 0.19 g/kg, n = 8) compared to 0.44 g/kg (SD = 0.12 g/kg, n = 11) in controls, t(17) = 2.21, *p* < 0.05 (Fig. 1E).

### Adult stress does not result in protracted alteration of ethanol self-administration

To determine whether the long-term effects of stress were specific to adolescent exposure, we examined long-term effects on ethanol self-administration in a group of rats treated with 14 days of CVS during adulthood starting on day PND 82 (Fig. 2A). Similar to the adolescent experiment described above, we waited at least 30 days after cessation of CVS to begin the self-administration procedures. Saccharin intake across the 2 days immediately prior to introduction of ethanol did not differ between adult control (Mean = 12.09 ml, SD = 2.09 ml, n = 9) and adult stress exposed rats (Mean = 10.6 ml, SD = 2.69 ml, n = 9), *p* > 0.05. In contrast to the effects of adolescent stress exposure, adult stress did not result in long-term changes in alcohol self-administration of saccharin sweetened ethanol (4.0% EtOH + 0.1% Sac) over 7 days, F(1, 16) = 0.55, *p* > 0.05; n = 9, or unsweetened ethanol (10% EtOH) across 10 days, F(1, 16) = 0.12, *p* > 0.05, n = 9 (Fig. 2B, D). The mean self-administered dose of ethanol across 8 days of saccharin sweetened 2-4% EtOH for adult control rats was 0.94 g/kg (SD = 0.14 g/kg, n = 9) and 0.91 g/kg, (SD = 0.13 g/kg, n = 9) for adult stress exposed rats, *p* > 0.05 (Fig. 2C). The mean dose across 10 days of 10% unsweetened ethanol was 0.49 g/kg (SD = 0.14 g/kg, n = 9) for controls and 0.44 g/kg (SD = 0.13 g/kg, n = 9) for adult stress exposed rats, *p* > 0.05 (Fig. 2E). These results indicate that adolescent, but not adult, exposure to CVS results in long-term increased operant self-administration of ethanol. Prior work from our lab found that adolescent chronic nicotine results in increased ethanol intake along with altered VTA GABA signaling 30 days after nicotine cessation, but adult nicotine exposure did not alter either (Thomas et al., 2018). Therefore, we next examined if adolescent stress resulted in long-term changes in VTA GABA signaling.

**Figure 2.**
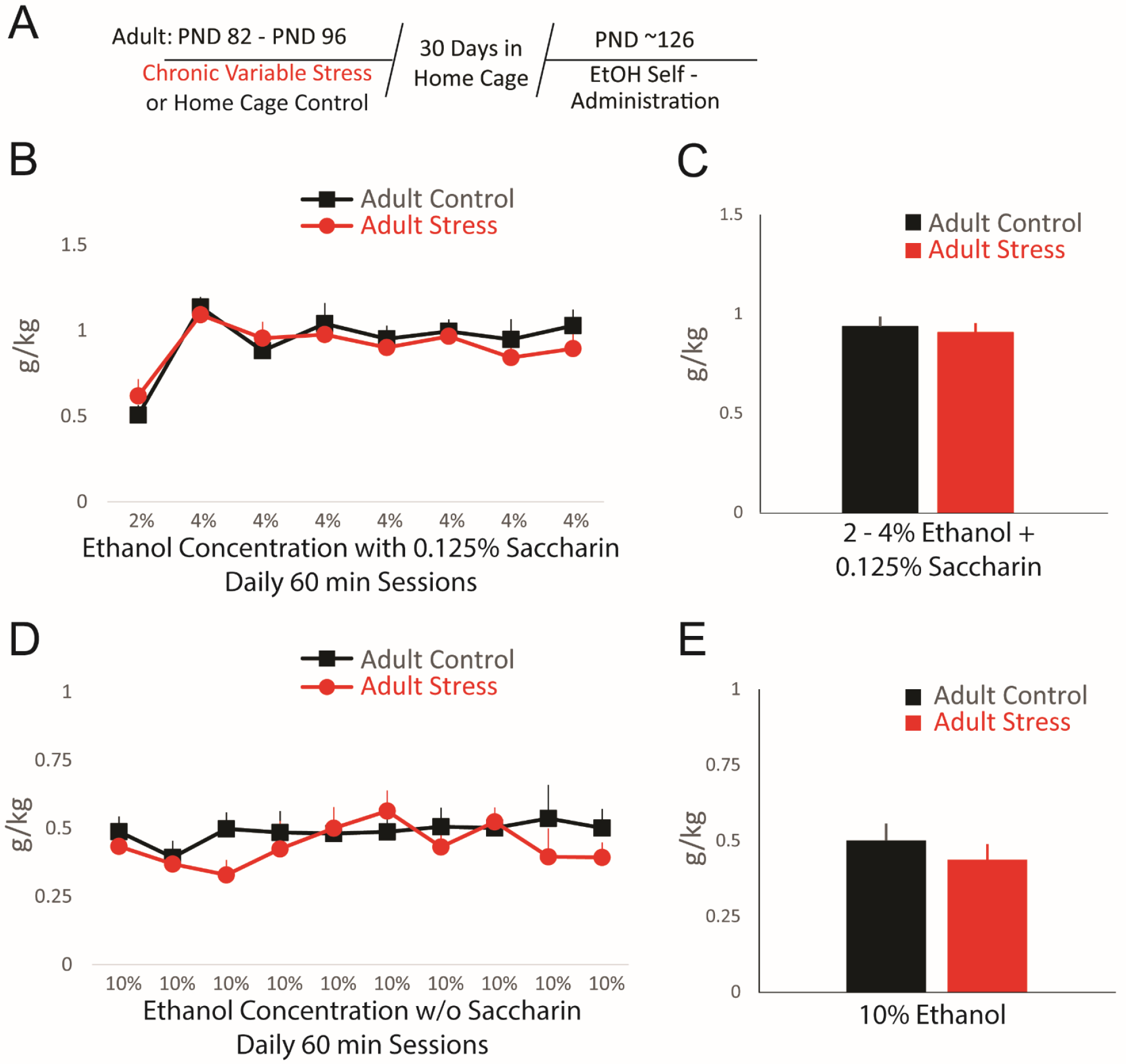
(A) Adult rats were exposed to chronic variable stress for 14 days, PND 82 - 96. Ethanol self-administration was assessed at least 30 days after cessation of stress. (B) Mean daily ethanol intake during initial self-administration sessions. Adult stress-exposed rats (red) showed do difference in daily intake of sweetened (0.1% saccharin) 4% ethanol compared to home cage controls (black). *p* > 0.05, mixed-design ANOVA, n = 9 rats per group. (C) Average ethanol self-administration of 2-4% ethanol with saccharin 0.1%. Adult stress exposed rats showed no difference in ethanol intake compared to controls. *p* > 0.05, independent samples t-test, n = 9 rats per group. (D) Mean daily intake of 10% ethanol after removal of saccharin over 10 self-administration sessions. Ethanol intake was not different in adult stress-treated rats compared to controls. *p* > 0.05, mixed-design ANOVA, n = 9 rats per group. (E) Average ethanol self-administration of unsweetened 10% ethanol over 10 sessions. Adult stress exposed rats showed no difference in 10% ethanol intake compared to controls. *p* > 0.05, independent samples t-test, n = 9 rats per group

### Adolescent chronic variable stress alters ethanol-induced VTA GABA transmission

Enhanced ethanol self-administration has been previously associated with the increased ethanol-induced GABAergic transmission onto VTA DA neurons (Ostroumov et al., 2016; Thomas et al., 2018). In order to determine whether adolescent stress exposure alters GABA transmission onto DA neurons in the VTA, we performed whole-cell recordings of VTA dopamine neurons and measured spontaneous IPSCs (sIPSC) in the presence of ethanol (Fig. 3A). In midbrain slices cut from unstressed control rats, we observed a modest increase in the sIPSC frequency following bath-applied ethanol (50 mM): 117.0% ± 5.6% of basal (Figures 3B and 3C, black data). However, in midbrain slices cut from stressed rats, we observed a significantly greater ethanol-induced potentiation of sIPSC frequency compared to control (Fig. 3B and 3C, red data): 158.1% ± 8.0% of basal, n = 8 cells, 4-5 animals per group, *p* < 0.01. There were no significant differences in sIPSC amplitude between the stress and control groups upon exposure to ethanol: 101.6 % ± 3.9 % in control rats and 104.3% ± 4.4% in stressed rats, *p* > 0.05, n = 8 cells, 4-5 animals per group. In addition, between groups there were no significant differences in baseline sIPSC frequency (3.72 ± 1.07 Hz among control rats versus 3.85 ± 0.67 Hz among stressed rats, *p* > 0.05) or baseline amplitude (29.5 ± 2.9 pA among control rats versus 35.2 ± 2.2 pA among stressed rats, *p* > 0.05). The significant increase in sIPSC frequency, but not amplitude, among stressed rats indicates an enhanced presynaptic release of GABA in the presence of ethanol.

**Figure 3.**
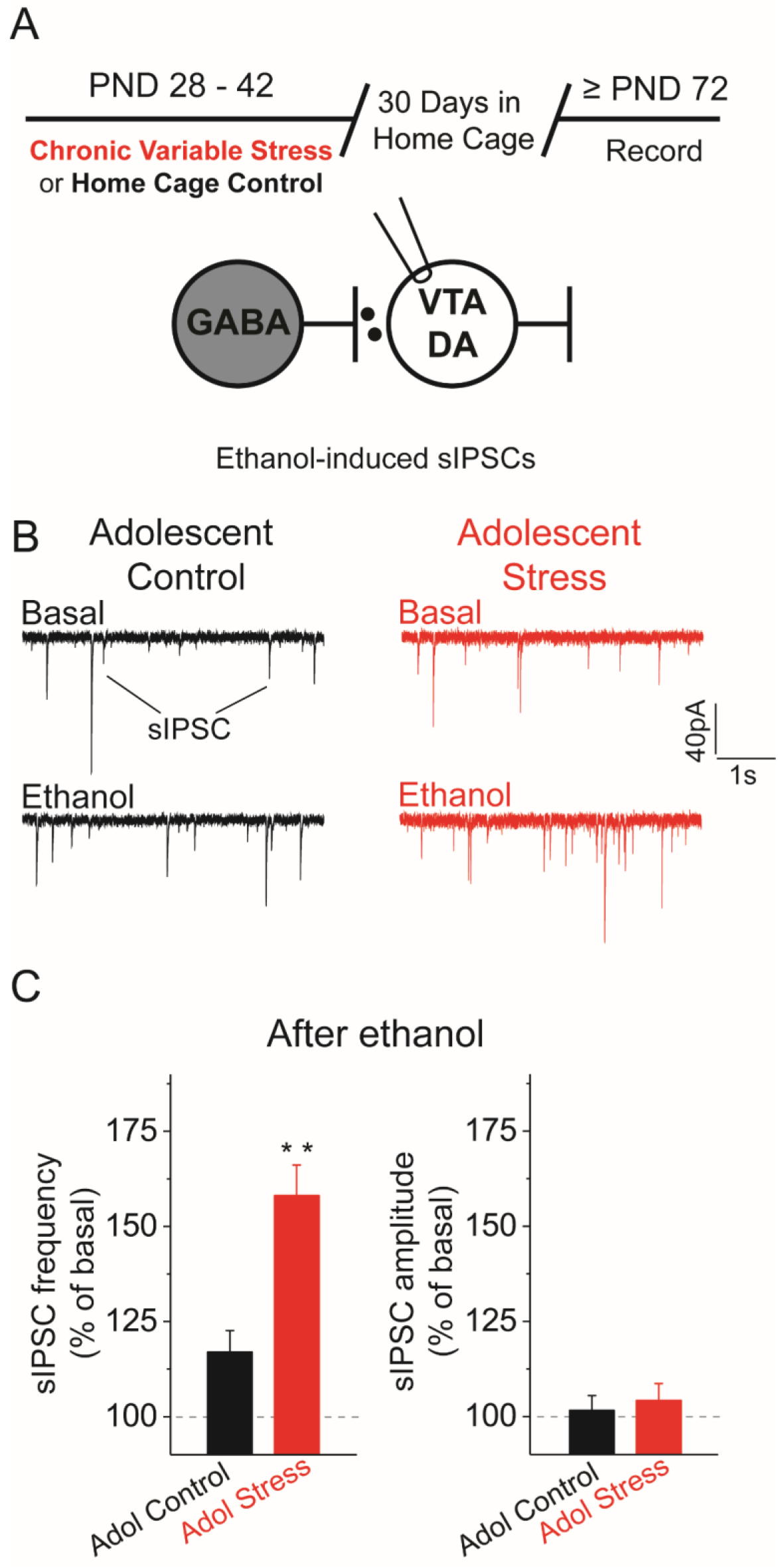
(A) The experimental time line is shown along with a didactic representation of spontaneous inhibitory postsynaptic currents (sIPSCs) onto VTA DA neurons, which were recorded from midbrain slices using the whole-cell patch-clamp configuration in the presence of glutamate receptor antagonists. (B) Representative recordings of sIPSCs before and after ethanol ex vivo application in control (black) and adolescent stress exposed (red) rats. No significant differences were observed in the mean basal sIPSC frequency or amplitude between stress-treated and control rats prior to ethanol. (C) Mean changes in the sIPSC frequency after bath application of ethanol (50 mM) in VTA DA neurons. DA neurons from stress exposed animals (red) demonstrated a significant increase in ethanol-mediated sIPSC frequency compared to neurons from controls. \*\**p* < 0.01, independent samples t test, n = 8 cells, 4-5 animals per group

### Adolescent chronic variable stress decreases chloride extrusion capacity in VTA GABA neurons

Enhancement of ethanol GABA transmission onto DA neurons may be due to increased excitability of VTA GABA neurons synapsing on VTA DA neurons. Our lab has previously shown that following acute adult stress VTA GABA neurons fail to maintain a normal Cl^−^ gradient, leading to a decrease in inhibitory signaling efficiency at GABA_A_ receptors on VTA GABA neurons (Ostroumov et al., 2016). To test if adolescent stress (Fig. 4A) exposure weakens Cl^−^ extrusion from VTA GABA neurons, we applied repetitive GABA_A_ receptor stimulation (25 pulses at 20 Hz) during whole-cell recordings (Fig. 4B) and measured activity-dependent depression of the evoked IPSCs. This experiment was performed at two holding potentials. First, the neuron was held at 0 mV, which prompts chloride influx into the cell through GABA_A_ receptors upon synaptic stimulation. A significantly greater depression in evoked IPSC amplitudes was observed in GABA neurons from stressed rats (red) as compared to control rats (black): F(2;5) = 25.57, *p* < 0.01, n = 8 cells, 4 animals per group (Fig. 4C and 4D). The rate of decrease in evoked IPSC amplitude at 0 mV is dependent upon intracellular chloride accumulation and consequent activity-dependent synaptic depression. To test the possibility that increased evoked IPSC amplitude depression in the stressed group was due to presynaptic rundown, GABA neurons were held at −90 mV, when the chloride driving force is outward. There were no differences in synaptic depression between stress and control groups at −90 mV: F(2;5) = 1.48, *p* > 0.05, n=8 cells, 4 animals per group (Fig. 4E and 4F). For this reason, the faster rates of IPSC amplitude reduction during repetitive stimulation at 0 mV are indicative of impaired chloride extrusion (Hewitt et al., 2009)(Ostroumov et al., 2016).

**Figure 4.**
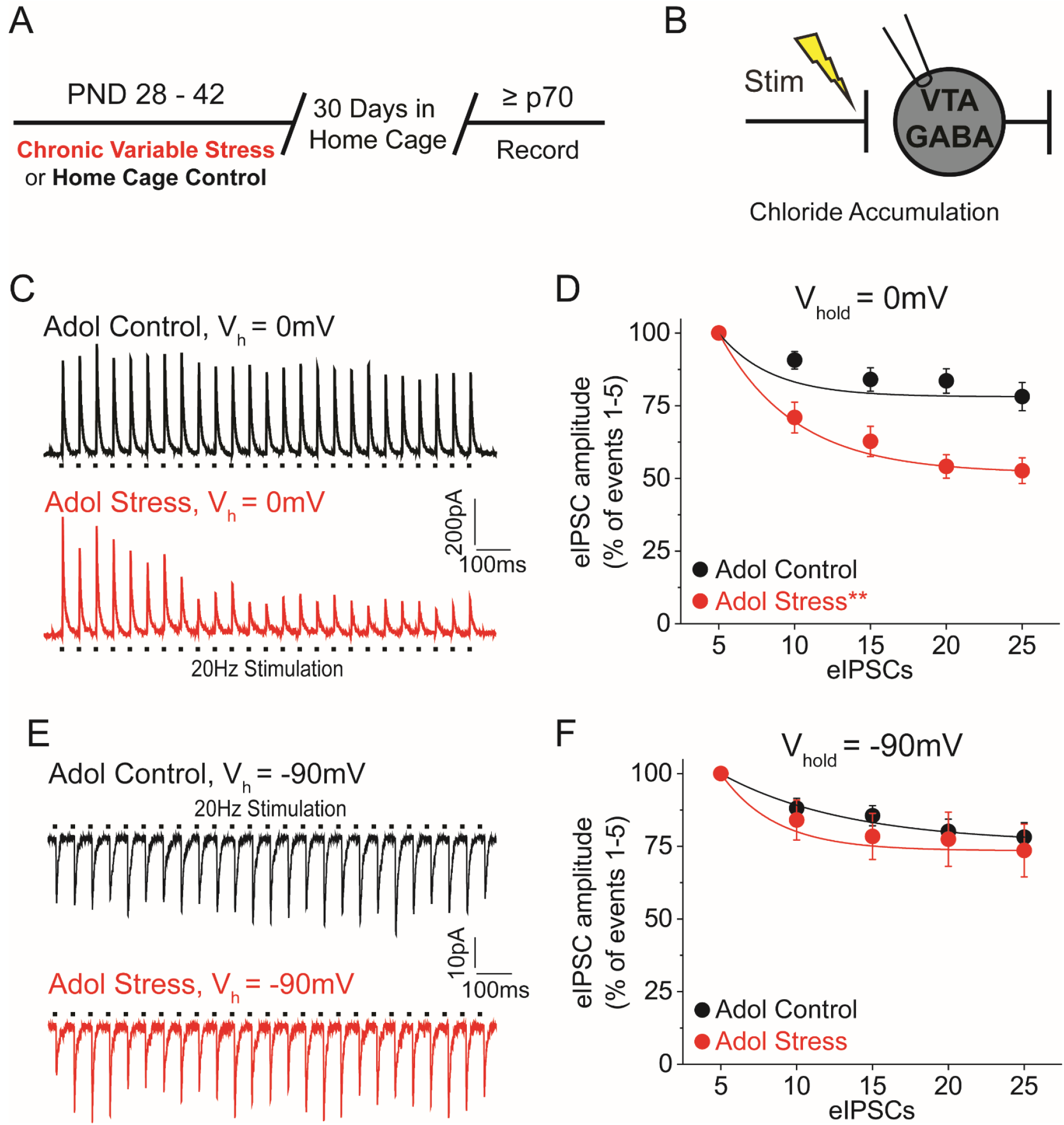
(A) Experimental timeline is shown. Animals were exposed to chronic variable stress for two weeks during adolescence, then recordings were performed at least 30 days later. (B) The effect of stress on Cl^−^ intracellular accumulation was estimated by repetitively stimulating GABA_A_R input and measuring the evoked IPSC amplitude from VTA GABA neurons. (C) Upon stimulation (25 pulses at 20 Hz, V_h_ = 0 mV causing Cl^−^ influx), a representative GABA neuron from a control animal (black) demonstrated a minor depression of IPSC amplitude compared to the significantly greater depression seen in a GABA neuron from an adolescent stress treated animal (red). (D) At 0 mV, VTA GABA neurons from adolescent stress-treated animals (red) demonstrated a significantly greater rate of evoked IPSC amplitude depression than GABA neurons from control animals (black) \*\**p* < 0.01, significantly different from the control by F test, 8 cells, 4 animals per group. (E) Activity dependent synaptic depression was examined during whole-cell patch clamp recordings under repetitive GABA_A_ receptor input. Holding at −90 mV (V_h_ = −90 mV causing Cl^−^ efflux), a representative GABA neuron from control (black) and stressed (red) rats showed a similar evoked IPSC amplitudes when stimulated at 20 Hz. (F) At −90 mV, VTA GABA neurons from control (black) and stressed (red) animals demonstrated no significant difference in rate of evoked IPSC amplitude depression, *p* > 0.05, n = 8 cells, 4 animals per group.

## Discussion

Early life stress and adversity predict heightened risk for AUDs and greater alcohol intake, but the underlying neurobiological adaptations that mediate this relationship between stress and ethanol are not well understood. In the present study, we identified age-dependent long-term effects of stress on ethanol self-administration in adult rats. Specifically, we found that rats exposed to CVS during adolescence (PND 28-42), then maintained in their home cages without stress until adulthood, had long-term increased self-administration of sweetened and unsweetened ethanol in adulthood. In contrast, rats that were exposed to CVS as adults (PND 82-96), and were given the same length of time in their home cages without stress, did not show long-term changes in alcohol self-administration. Concomitant with adolescent stress-induced increased ethanol self-administration in adulthood, we found increased ethanol-induced inhibitory signaling onto VTA DA neurons. We also observed later life deficits in chloride extrusion in VTA GABA neurons of rats exposed to adolescent stress (Ostroumov et al., 2016).

Adolescence demarks a period of critical neural development, during which the individuals are at increased vulnerability to long-term consequences of stress (Spear, 2000). Our finding, that adolescent, but not adult, stress resulted in a long-term increase in ethanol self-administration is in agreement with prior work examining the effects of adolescent social isolation stress (Butler et al., 2016; Lopez et al., 2011; Schenk, Gorman, & Amit, 1990; Skelly, Chappell, Carter, & Weiner, 2015). Similarly, age-dependent stress effects have been observed in neurobiological and stress function (Cotella et al., 2019; Yorgason, Espana, Konstantopoulos, Weiner, & Jones, 2013). Importantly, our results are in agreement with a prior study that implemented adolescent CVS and found increased adult ethanol intake using 2 bottle choice (Lopez et al., 2011). One important difference in our study was the use of operant self-administration, which allowed for rats to be group housed throughout the ethanol intake testing period (except for the 1-hr self-administration session). Because single housing is a known stressor, our findings suggest that the effects of adolescent stress on alcohol intake behaviors is independent of proximal or concurrent social isolation stress. Additionally, stress effects on ethanol intake are dependent to some degree on the method of measurement (two-bottle choice vs. operant self-administration) (Noori, Helinski, & Spanagel, 2014). Therefore, our positive results using operant self-administration indicate that adolescent CVS is a robust model of early life stress.

We found that adolescent stress resulted in increased ethanol-induced inhibition of VTA DA neurons. These findings suggest that adolescent stress can result in long-term adaptations in VTA circuit function. A prior study found adolescent rats exposed to combined foot shock and restraint stress had increased DA excitability within the lateral VTA (Gomes & Grace, 2017). Interestingly, these results were stress type-dependent, and restraint and foot shock alone did not result in changes in VTA activity. Indeed, the effects of stress appear sensitive to the stress paradigm, and further work is required to parse out fundamental aspects of the particular stress exposures (Noori et al., 2014). It also has been shown that drugs of abuse, including ethanol, potently modulate GABA and DA signaling within the VTA (Nestler, 2005). Prior work from our lab demonstrated that acute stress potentiates ethanol-induced VTA GABA activity in both ex vivo and in vivo preparations (Ostroumov et al., 2016). Moreover, in midbrain slices, we have shown that acute stress increases ethanol-induced GABA signaling onto VTA DA neurons (Ostroumov et al., 2016). Therefore, our data suggest that when stress exposure occurs during adolescence the effect of stress on VTA GABA transmission onto DA neurons is long-lasting. Additionally, we observed long-term deficits in VTA GABA Cl^−^ extrusion, suggesting that the increased excitability of these cells onto the DA neurons may be due to an accumulation of intracellular chloride. Consequently, afferent GABAergic activity onto local VTA GABA neurons would not be as inhibitory and could become paradoxically excitatory (Ostroumov et al., 2016; Thomas et al., 2018).

Previously, in adult animals, stress and drugs of abuse were shown to cause a depolarizing shift in GABA_A_ receptor signaling within the VTA in concert with a deficit in the chloride transport mechanism of VTA GABA neurons (Ostroumov et al., 2016; Taylor et al., 2016; Thomas et al., 2018). Similarly, adolescent nicotine administration was found to result in long-term depolarizing shifts in GABA_A_ receptor signaling in adulthood (Thomas et al., 2018). Adult acute proximal stress and adolescent nicotine-induced depolarizing shifts in GABA_A_ signaling were shown to be due to decreases in the function of the K^+^, Cl^−^ transporter, KCC2, which is critical for maintenance of Cl^−^ homeostasis (Furukawa et al., 2017; Ostroumov et al., 2016; Plotkin, Snyder, Hebert, & Delpire, 1997). When KCC2 was functionally upregulated pharmacologically, ethanol self-administration was decreased in adult acute-stress exposed rats (Ostroumov et al., 2016). Therefore, persistent deficits in Cl^−^ extrusion in VTA GABA cells after adolescent stress may be the result of downregulated KCC2 functioning. Furthermore, our data are parsimonious with prior work demonstrating that increased GABA excitability due to decreased Cl^−^ extrusion capacity in VTA GABA neurons drives increased inhibitory signaling onto DA neurons in the presence of ethanol (Ostroumov & Dani, 2018a, 2018b; Ostroumov et al., 2016; Thomas et al., 2018).

In summary, adolescent, but not adult, stress results in long-term increases in operant ethanol self-administration and adaptations of midbrain GABA signaling. Our work is in agreement with other age-specific behavioral and neurophysiological effects, demonstrating that adolescence is a time of heightened vulnerability to the long-term consequences of a range of perturbations, including stress and drugs of abuse (Chin, Van Skike, & Matthews, 2010; Connor & Gould, 2017; Lopez et al., 2011; Thomas et al., 2018). Moreover, Cl^−^ homeostasis in VTA GABA neurons is vulnerable to adolescent stress, and this decreased Cl^−^ cellular gradient critically alters inhibition of VTA DA neurons in response to ethanol. Finally, our results add to a developing literature suggesting that stress-dependent potentiation of ethanol consumption might be combated with treatments that reverse Cl^−^ extrusion associated adaptations in GABA circuitry within stress and trauma exposed populations (Ostroumov & Dani, 2018b; Ostroumov et al., 2016; Thomas et al., 2018).

## Acknowledgments

The authors would like to acknowledge grant support from the National Institute on Drug Abuse (R01DA009411), to J.A.D. and postdoctoral fellowship from the National Institutes of Mental Health (2T32MH014654-40), to D.C.

